# *In vivo* Chemical Reprogramming of Astrocytes into Functional Neurons

**DOI:** 10.1101/305185

**Authors:** Yantao Ma, Handan Xie, Xiaomin Du, Lipeng Wang, Xueqin Jin, Shicheng Sun, Yanchuang Han, Yawen Han, Jun Xu, Zhuo Huang, Zhen Chai, Hongkui Dengi

**Affiliations:** Department of Cell Biology, School of Basic Medical Sciences, Peking University Stem Cell Research Center, State Key Laboratory of Natural and Biomimetic Drugs, Peking University Health Science Center, Beijing 100191, China; MOE Key Laboratory of Cell Proliferation and Differentiation, College of Life Sciences, Peking-Tsinghua Center for Life Sciences, Peking University, Beijing 100871, China.; Shenzhen Stem Cell Engineering Laboratory, Key Laboratory of Chemical Genomics, Peking University Shenzhen Graduate School, Shenzhen, Guangdong 518055, China.; Academy for Advanced Interdisciplinary Studies, Peking University, Beijing 100871, China. Peking University-Tsinghua University-National Institute of Biological Science Joint Gradute Program, College of Life Science, Peking University, Bejing 100871, China.; Peking University-Tsinghua University-National Institute of Biological Science Joint Gradute Program, College of Life Science, Peking University, Bejing 100871, China.; Department of Molecular and Celluar Pharmacology, School of Pharmaceutical Sciences, Health Science Center of Peking University, Bejing 100871, China.; State Key Laboratory of Membrane Biology, College of Life Sciences, Peking University, Beijing 100871, China.

## Abstract

Mammals lack robust regenerative abilities. Lost cells in impaired tissue could potentially be compensated by converting nearby cells *in situ* through *in vivo* reprogramming. Small molecule-induced reprogramming is a spatiotemporally flexible and non-integrative strategy for altering cell fate, which is, in principle, favorable for the *in vivo* reprogramming in organs with poor regenerative abilities, such as the brain. Here, we demonstrate that in the adult mouse brain, small molecules can reprogram resident astrocytes into functional neurons. The *in situ* chemically induced neurons (CiNs) resemble endogenous neurons in terms of neuron-specific marker expression and electrophysiological properties. Importantly, these CiNs can integrate into the mouse brain. Our study, for the first time, demonstrates *in vivo* chemical reprogramming in the adult brain, which could be a novel path for generating desired cells *in situ* for regenerative medicine.

---

The fate of committed cells can be reprogrammed by cellular, genetic and chemical approaches, which offers the possibility of generating desired cells by cell fate conversion (*1–5*). Remarkably, advances in chemical approaches have revealed that somatic cells can be reprogrammed into pluripotent stem cells completely by small molecules (*5*), and importantly, chemical reprograming is capable of inducing direct lineage conversion between differentiated cell types (*6–10*). Small molecules are easily manipulated and flexibly combined to reprogram cell identities at both the temporal and spatial levels to generate functional cells (*11, 12*). Furthermore, chemical approaches are cell permeable, reversible and non-integrative to the genome, which is favorable for *in vivo* reprogramming. Chemically induced *in vivo* reprogramming could, in principle, produce functional cells by converting nearby cells *in situ* to compensate for the cellular loss in damaged organs with limited regenerative capability, such as those of the central nervous systems (CNS). The mammalian brain does not readily repair damage because the loss of non-proliferative neurons is irreversible, and neural stem cells can only give rise to neurons in few brain regions. However, upon brain injury, neighboring astrocytes can become proliferative and serve as an abundant resource for reprogramming (*13*). For these reasons, our study aims to chemically reprogram local astrocytes in the adult brain, which could potentially regenerate compensatory neurons *in situ.*

Our previous chemically defined cocktail FICB (<u>F</u>orskolin, <u>I</u>SX9, <u>C</u>HIR99021 and I-<u>B</u>ET151) could reprogram fibroblasts into neuron-like cells *in vitro* (*7*). We therefore tested this cocktail on astrocytes carrying neuron-specific TauEGFP reporter for neuronal induction (*14*) (fig. S1). Upon exposure to FICB, astrocytes underwent neuron-like morphological changes with neurite outreach, and TauEGFP^+^ cells were observed at 16 day-post-induction (dpi), suggesting the acquisition of neuronal cell fate. An optimized cocktail DFICBY (<u>d</u>bcAMP, <u>F</u>orskolin, <u>I</u>SX9, <u>C</u>HIR99021, I-<u>B</u>ET151 and <u>Y</u>-27632) enhanced neuronal conversion efficiency, yielding 89.2 ± 1.4 % of TUJ1^+^ cells with a neuron-like morphology at 16 dpi (fig. S1, B and C), which cells were immunopositive for the neuron-specific markers MAP2, SYN1, and NF-H (fig. S1B). The mature neuronal marker NEUN was detected at 30 dpi with an efficiency of 77.8 ± 11.1 %, when these cells were co-cultured with primary astrocytes from 16 dpi in a defined maturation medium FCBG (<u>F</u>orskolin, <u>C</u>HIR99021, brain derived neurotrophic factor (<u>B</u>DNF) and glial cell line-derived neurotrophic factor (<u>G</u>DNF)) for an additional two weeks. Meanwhile, the glutamatergic neuronal marker vGLUT2 and GABAergic neuronal marker GAD67 were also detected, suggesting that both excitatory and inhibitory neurons were generated (fig. S1, B and E). Wholecell recording showed that the CiNs were capable of generating action potentials (APs) in response to depolarizing step currents (fig. S1F, n = 12) and inactivating inward currents (fig. S1F; Table S1; n = 11). We also confirmed the astroglial origin of CiNs with lineage-tracing mice carrying *mGfap-Cre/Rosa26*-tdTomato/TauEGFP (*15, 16*) (fig. S2, A and B). Nearly all (96.5 ± 1.1 %) tdTOMATO^+^ cells were immunopositive for astroglia-specific marker (S100B) and negative for neural markers (NEUN, DCX), microglial marker (IBA1) and oligodendrocyte marker (O4) (fig. S2, B and C). At 16 dpi with DFICBY treatment, we observed that 97.4 ± 1.2 % of TUJ1^+^ cells co-expressed tdTOMATO, suggesting that nearly all CiNs were converted from astrocytes (fig. S2D). Collectively, DFICBY, that robustly reprogrammed astrocytes into CiNs with functional neuronal properties.

Next, we tested whether the chemical cocktail could also induce CiNs from astrocytes *in situ* in adult mouse brains. To trace the astroglial origin of CiNs *in vivo*, we generated lineage-tracing mice by crossing *mGfap-Cre* mice with *Rosa26*-tdTomato reporter mice (Fig. 1A). In the crossed mice, tdTOMATO^+^ cells in the striatum co-expressed the astrocyte markers GFAP (95.6 ± 4.0 %) and S100B (91.3 ± 7.0 %) and were negative for microglia marker IBA1 and oligodendrocyte marker MBP, suggesting a high specificity of the lineage-tracing system (Fig. 1, B and C). Rare tdTOMATO^+^ cells (0.8 ± 0.2 %) expressed the neuronal cell marker NEUN but not proneural markers DCX, NEUROD1 and NEUROG2 (Fig. 1, B and C). Thus, initiating astrocytes for the *in vivo* reprogramming in the striatum of adult mouse brain were specifically labeled by tdTOMATO in *mGfap-Cre/Rosa26*-tdTomato mice.

**Figure 1.**
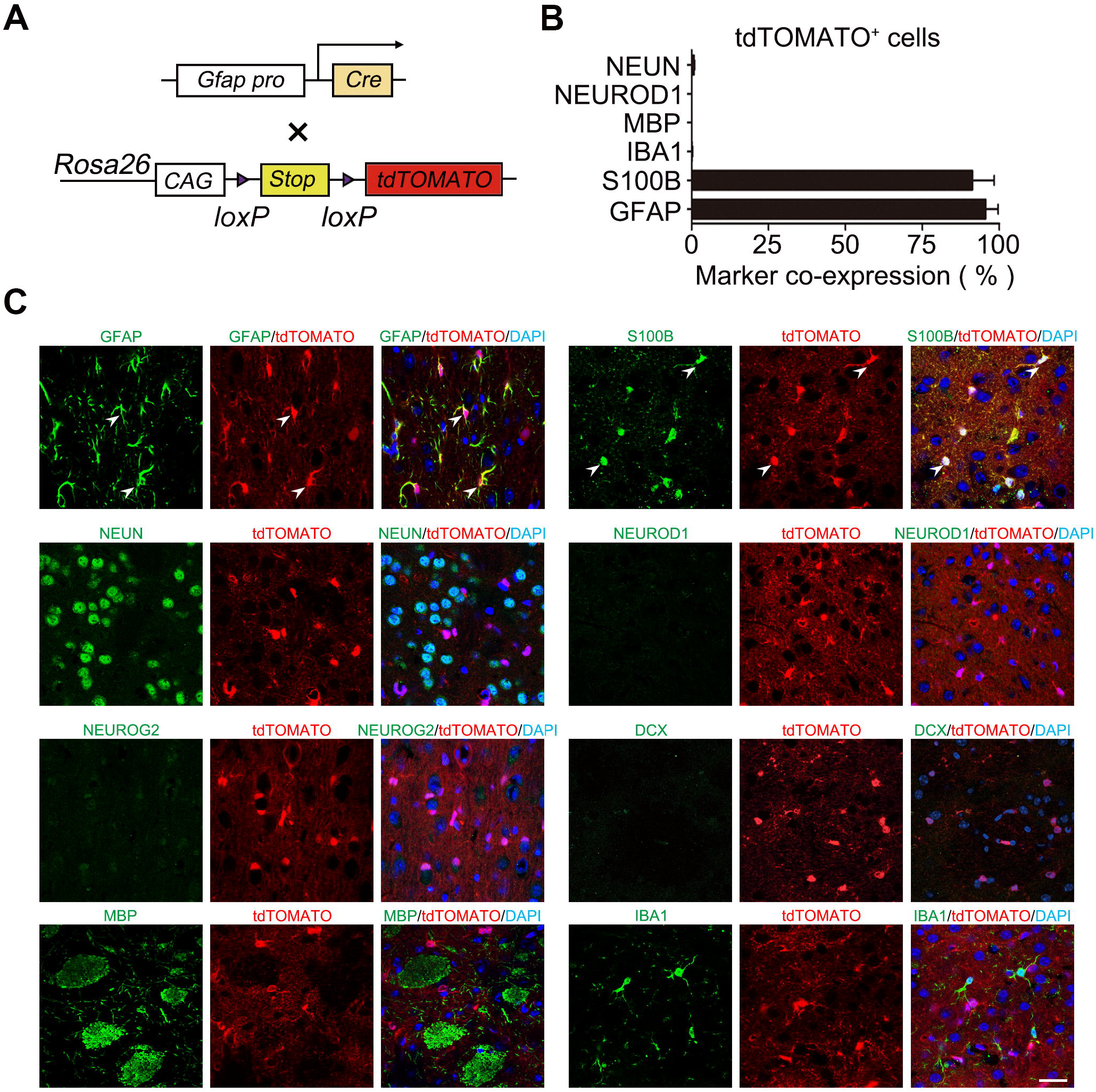
Lineage tracing system to track the astrocytic origin of converted CiNs. (A) Scheme of *mGfap-Cre/Rosa26*-tdTomato lineage-tracing constructs. (B) Cell identity characterization of tdTOMATO^+^ cells in lineage-tracing mice (n = 3). (C) Immunohistochemistry characterization of tdTOMATO^+^ cells in lineage-tracing mice. tdTOMATO^+^ cells were immunopositive for astroglial markers GFAP and S100B. These cells were negative for neuronal markers (NEUN, DCX, NEUROD1 and NEUROG2), microglial marker (IBA1) and oligodendrocyte marker (MBP). Arrowheads point to examples of tdTOMATO labeled cells. Scale bar: 25 μm. Error bars represent s. e. m.

To induce chemical reprogramming *in vivo*, the small molecule cocktail was administered into the striatum of 8-week old mice at a constant rate for two weeks via an osmotic mini-pumping system (Fig. 2A). After DFICBY induction, we observed tdTOMATO^+^/NEUN^+^ cells in the striatum at 8 week-post-injection (wpi), suggesting that this cocktail could reprogram astrocytes into neuron-like cells *in vivo* (data not shown). We further optimized the DFICBY cocktail for *in vivo* reprogramming and found that partial substitution of dbcAMP for FSK did not overcome FSK to the efficiency of *in vivo* conversion. Additionally, we found that increasing the dose of FICBY by up to threefold in the *in vitro* dose led to a higher efficacy *in vivo* (fig. S3A). We next analyzed the effect of individual small molecules on the *in vivo* neuronal reprogramming. By single small molecule omission, we found that removing Forskolin, I-BET or ISX9 greatly hindered the generation of CiNs *in vivo*; and in the absence of CHIR99021, we observed a significant reduction in reprogramming efficiency (fig. S3B), suggesting that small molecules reprogrammed astrocytes to CiNs *in vivo* in a synergetic manner. At 8 wpi, the optimized formula FICBY generated tdTOMATO^+^/NEUN^+^ cells in the striatum (Fig. 2B).

**Figure 2.**
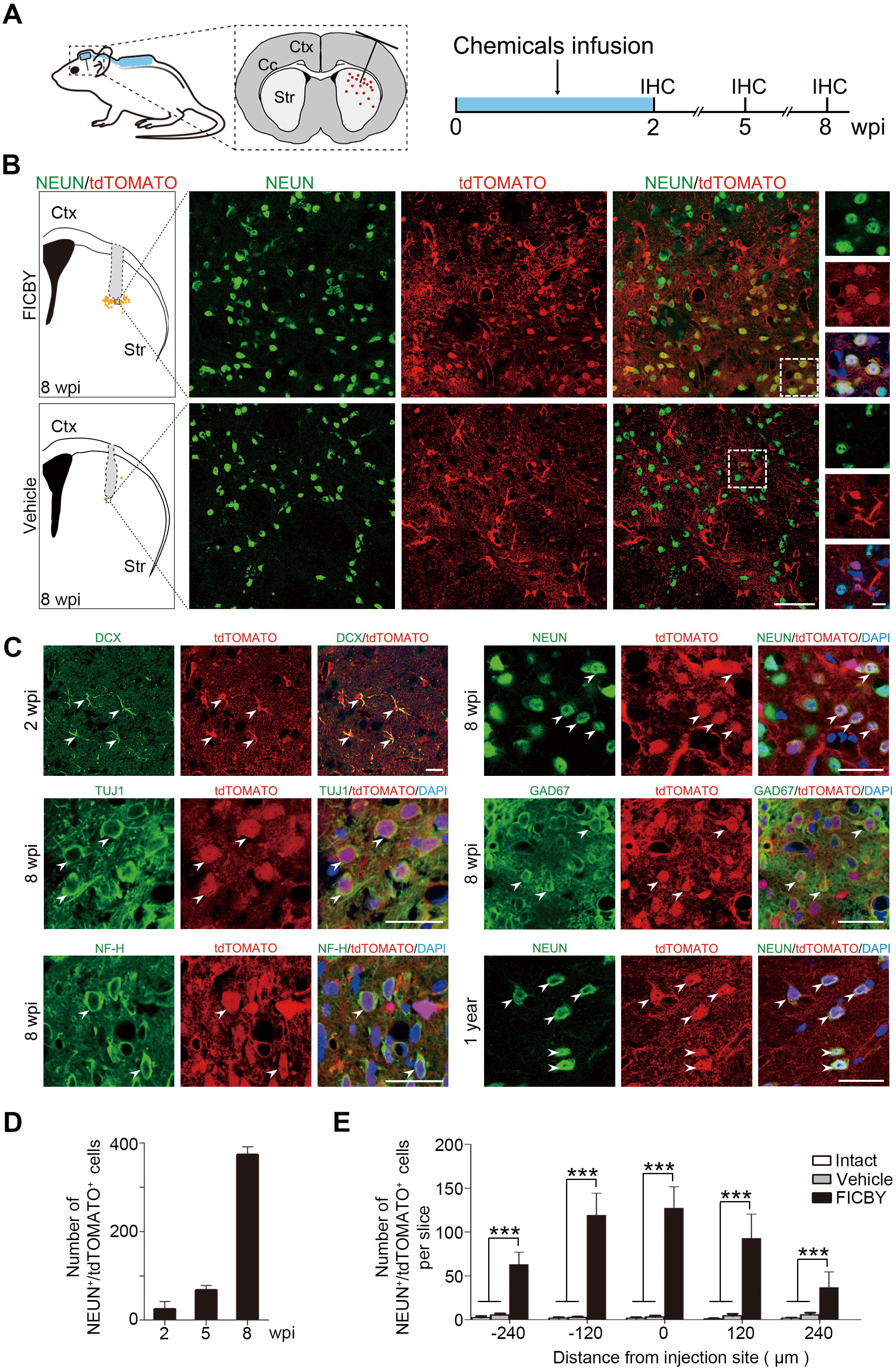
*In vivo* chemical induction of resident astrocytes into CiNs in adult mouse striatum. (A) Scheme of the chemical infusion into the mouse brain. (IHC, immunohistochemistry; Ctx, cortex; Cc, corpus callosum; Str, striatum) (B) Diagram of the distribution of converted neurons in the striatum. Immunohistochemistry analyses of NEUN^+^/tdTOMATO^+^ cells in the striatum of chemically treated mice at 8 wpi. High-magnification panels represent the co-localization of NEUN and DAPI. Yellow dots represent NEUN^+^/tdTOMATO^+^ cells in the striatum. (C) Immunohistochemistry analyses of CiNs at different time points; in tdTOMATO^+^ cells, the DCX were detected at 2 wpi; TUJ1, NF-H and GAD67 were detected at 8 wpi. NEUN were detected at both 8 wpi and one year-post-injection. Arrowheads point to examples of CiNs. (D) Quantifications of NEUN^+^/tdTOMATO^+^ cells around the injection site in the striatum of chemically treated mice at 2, 5, and 8 wpi (n = 2). (E) Quantification of NEUN^+^/tdTOMATO^+^ cells at different distances from the injection site at 8 wpi. (intact group, n = 3; vehicle group, n = 5 and FICBY group, n = 5). Scale bars: (B) 75 μm, 20 μm in high-magnification panels; (C) 25 μm. Error bars represent s. e. m. *** *P* < 0.0001 by one-way ANOVA with Tukey’s Multiple Comparison Test.

During this reprogramming process, at 2 wpi, the tdTOMATO^+^ cells were immunopositive for the immature neuronal maker DCX and showed a mixed astrocyte-to-neuron morphology (Fig. 2C). At 5 wpi, tdTOMATO^+^/NEUN^+^ cells were detected. The number of NEUN^+^ CiNs increased from 5 to 8 wpi, suggesting gradual maturation of the CiNs after chemical induction (Fig. 2D). At 8 wpi, the tdTOMATO^+^ cells also expressed other neuronal-specific markers including TUJ1 and NF-H (Fig. 2C). Interestingly, tdTOMATO^+^ cells co-expressing the neuronal subtype marker GAD67 were also detected, suggesting that the small molecules could induce inhibitory CiNs from astrocytes *in vivo* (Fig. 2C).

To systemically quantify CiNs distribution, we analyzed 5 brain slices per mouse throughout the rostral-caudal extent on either side of the cannula at 8 wpi, which was considered the main distribution region of the small molecules (Fig. 2E). Scattered CiNs were distributed locally throughout the outer layer of the injection site, suggesting a restrained effective area of small molecules in the brain (Fig. 2B). Notably, 127 ± 24 tdTOMATO^+^/NEUN^+^ cells per slice (n = 5) were observed at the injection core, while only a few tdTOMATO^+^/NEUN^+^ cells were observed in the vehicle group (3 ± 1 tdTOMATO^+^/NEUN^+^ cells, n = 5) and intact group (2 ± 1 tdTOMATO^+^/NEUN^+^ cells, n = 3) (Fig. 2E). NEUN^+^/tdTOMATO^+^ cell number did not show significant difference between intact group and vehicle group, which indicates surgery operation did not induce astrocyte-to-neuron cell fate conversion *in situ* (Fig. 2E). Furthermore, we observed tdTOMATO^+^/NEUN^+^ CiNs one year-post-injection, suggesting its long-term survival capacity (Fig. 2C). Notably, no solid brain tumors were observed in any of the tested mice.

Next, we tested whether the *in situ*-generated CiNs displayed electrophysiological functions. We performed dynamic analyses of brain slice recordings on tdTOMATO^+^ cells in the striatum with neuronal morphology around the injection site (Fig. 3A). Initially, at 0 wpi we randomly patched astrocytes on brain slices of reporter mice and found not competent for generating AP (Fig. 3C). Interestingly at 4 wpi, we observed membrane depolarization subjected to input current but still unable to generate AP, suggesting a transient state during the conversion toward neuronal electrophysiological property (Fig. 3B). AP of CiNs appeared as early as 6 wpi, followed by continuous APs at 8 wpi with a resting membrane potential of −74.5 ± 2.6 mV, an amplitude of 102.2 ± 3.5 mV and a threshold of −41.7 ± 5.3 mV (Fig. 3C; Table S2; n = 14). The percentage of CiNs which could fire more than two spikes increased along time (Fig. 3D). From 4 to 12 wpi, we observed the gradual maturation of membrane-intrinsic properties, as indicated by increased membrane capacitance (Cm) and a decrease in input resistance and AP threshold, which suggested that the CiNs resembled endogenous neurons over time (Fig. 3E; Table S2, 3). In addition, fast, inactivating inward currents were recorded in the CiNs at 8 wpi, which corresponds with the opening of voltage-dependent Na^+^ channels (Fig. 3F; n = 3). We further detected spontaneous excitatory post-synaptic currents (EPSCs) in the CiNs by blocking the inhibitory inputs, which was subsequently blocked by 6-cyano-7-nitroquinoxaline-2,3-dione (CNQX) and 2-amino-5-phospho-novaleric acid (AP5), resulting in the complete disappearance of EPSCs (Fig. 3G; n = 3). Collectively, these results indicated that in the mouse brain, CiNs acquired electrophysiological functions and formed synaptic connections.

**Figure 3.**
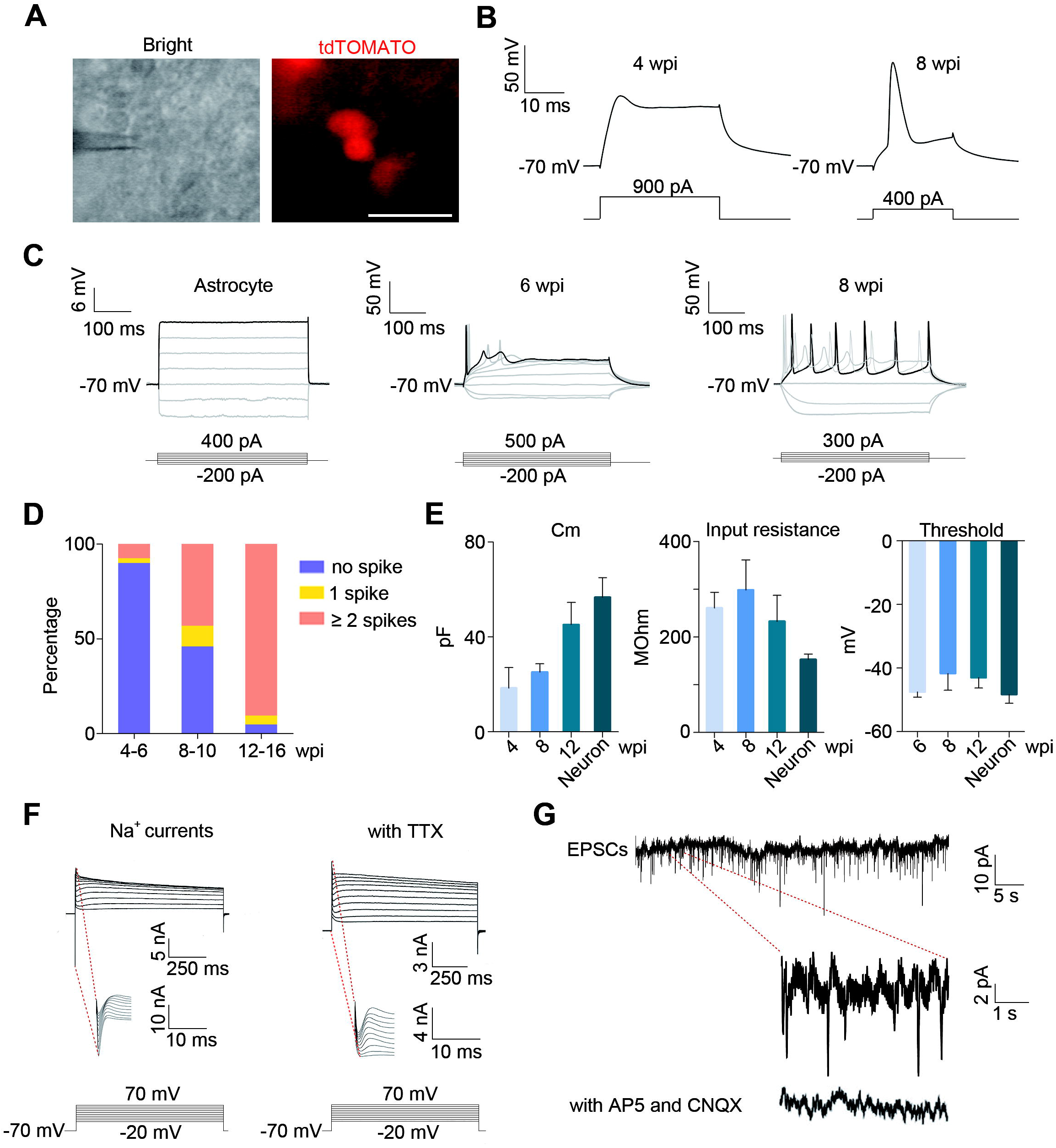
Electrophysiological analyses of *in vivo* reprogrammed CiNs. (A) Representative images of neuron-like tdTOMATO^+^ cells in brain slices forpatch-clamp recordings. (B) Representative single action potential (AP) traces of the CiNs at 4 wpi and 8 wpi. (C) Whole cell patch-clamp recordings of astrocytes, and the CiNs at 6 wpi and 8 wpi. (D) Quantification of CiNs firing none, one or more than one spike at various time points during *in vivo* chemical induction. (E) Gradual maturation of membrane-intrinsic properties of CiNs from 4 to 12 wpi as shown for membrane capacitance (Cm), input resistance and AP threshold. (F) Recording of inward currents in CiNs in voltage-clamp mode. Tetrodotoxin (TTX) (500 nM) was used to block voltage-dependent Na^+^ channels (n = 3). (G) Recording of spontaneous postsynaptic potentials in CiNs. CNQX (20 μM) and AP5 (50 μM) were used to block spontaneous postsynaptic potentials (n = 4). Scale bar: 20 μm. Error bars represent s. e. m.

Because the integration of compensatory neurons into the host neural circuitry is crucial for functional rescue (*17*), we further examined the connectivity of CiNs by a retrograde transsynaptic tracing system (*18*). We used a pseudotyped rabies virus (PRV) that specifically infected neurons expressing avian tumor virus receptor A (TVA). Subsequently, PRV assembled into infectious particles to cross one synapse where the rabies virus glycoprotein (RVG) was present. To achieve this, two Cre-dependent FLEX-AAV vectors encoding TVA, histone-EGFP and RVG were injected into the cortex of *mGfap-Cre* mice, resulting in their expressions restricted in astrocytes (Fig. 4A and fig. S4-5). After chemical induction, we injected PRV-carrying DsRed (PRV-DsRed) at 7 wpi into the same site to label the CiNs as DsRed^+^/EGFP^+^ cells and to trace endogenous host neurons that made synaptic contact with CiNs as DsRed single-positive cells (Fig. 4A). Besides, we found that after TVA injection, cortical astrocytes were specifically labeled while with minor specificity within the corpus callosum of the striatum. Therefore, we did transsynaptic experiments in the cortex.

**Figure 4.**
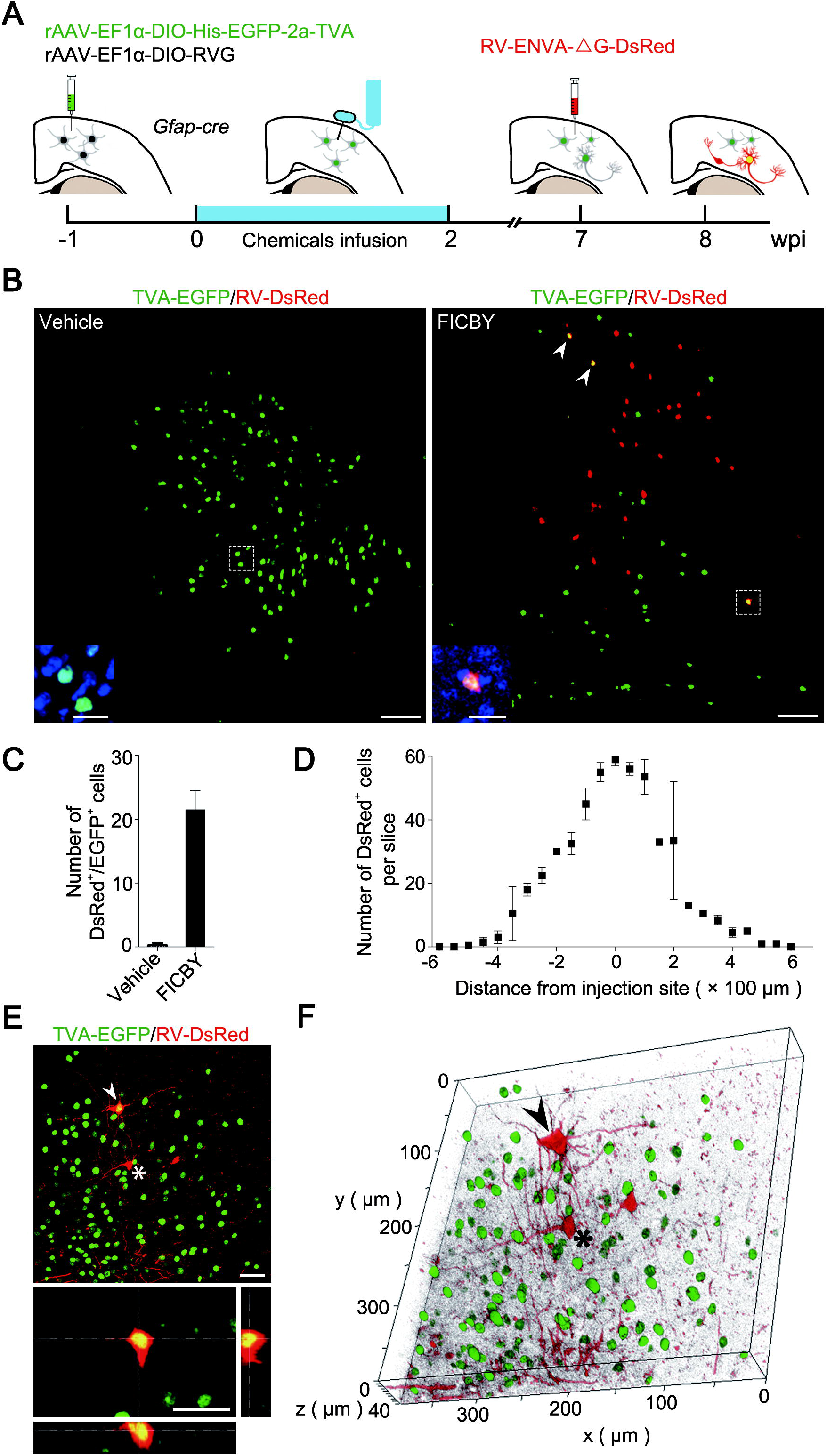
Integration of *in vivo* reprogrammed CiNs into the mouse brain. (A) Scheme of the PRV-based retrograde transsynaptic tracing process. (B) Confocal images of initiating astrocytes (green), CiNs (red and green) and traced host neurons (red) in the chemical induction group and vehicle group (n = 3). Arrowheads point to examples of CiNs. High-magnification panels represent the co-localization of EGFP and DAPI. (C) Histogram representing the total number of DsRed^+^/EGFP^+^ cells (CiNs) in the chemical induction group and vehicle group (n = 4). (D) Quantification of DsRed single-positive cells through the injection region (n = 2). (E) Representative confocal image of an EGFP^+^/DsRed^+^ CiN and its connecting DsRed single-positive neuron. White arrowhead points to the CiN, and the asterisk highlight the local neuron which has direct synaptic connections with the CiN. (F) The three-dimensional reconstruction of (E upper). The arrowhead and asterisk point to the same cells in (E upper). Scale bars: (B), 100 μm, 20 μm in high-magnification panels; (E), 40 μm. Error bars represent s. e. m.

At 8 wpi, we observed DsRed^+^/EGFP^+^ CiNs (22 ± 2 cells in total) at the injection site in the cortex of the chemical induction group (Fig. 4, B and C). Meanwhile, DsRed single-positive cells were detected in the area surrounding the injection site (Fig. 4, B and D). The presence of both DsRed^+^/EGFP^+^ and DsRed single-positive cells in the chemical treated group indicated that the CiNs were able to be innervated by host neurons (Fig. 4, E and F). In contrast, there were no DsRed^+^/EGFP^+^ cells observed in the vehicle group (Fig. 4C, n = 4). We also observed direct neurite connections between DsRed^+^/EGFP^+^ CiNs and DsRed single-positive cells, suggesting that synapses formed between CiNs and host neurons (Fig. 4, E and F). These data suggest that *in situ* converted CiNs could timely receive synaptic signals from the local neurons in the adult brain.

Our study is the first to demonstrate that defined combinations of small molecules can induce the *in vivo* chemical reprogramming of astrocytes into functional mature neurons with electrophysiological characteristics. Importantly, these *in situ*-generated CiNs could functionally interact with resident neurons in the brain. Astrocytes serve as a latent plastic cell type for neurogenesis (*19–21*), and our work suggests a new strategy of *in situ* chemical reprogramming for the endogenous repair of brain injuries and neurodegenerative diseases. Future studies with disease models are needed to investigate the neuronal subtype-specific functions of these CiNs and to develop safety chemical delivery systems to different regions of the CNS. *In vivo* chemical reprogramming avoids the shortcomings of cell transplantation, including immunorejection, stem cell mutagenesis, inadequate cell survival and native tissue integration difficulties (*22, 23*). Moreover, small molecule-induced *in situ* lineage conversion is non-immunogenic and transgene free, which is eventually preferable for *in vivo* therapeutic strategies(*24–28*). Our finding may also offer a general strategy for *in vivo* chemical reprogramming to produce cells of other lineages. *In vivo* chemical reprogramming would open a novel path for regenerative medicine.

## Acknowledgments

We would like to thank J.L. Wang, J.Y. Guan, Lin Cheng, M.L. Qu, Xu Zhang, Ting Zhao and other laboratory members for suggestions. This work was supported by the National Key Research and Development Program of China (2016YFA0100100 and 2017YFA0103000), the National Natural Science Foundation of China (31730059 and 31521004), the Guangdong Innovative and Entrepreneurial Research Team Program (2014ZT05S216), the Science and Technology Planning Project of Guangdong Province, China (2014B020226001), the Science and Technology Program of Guangzhou, China (2016B030232001), the Science and Technology Program of Guangzhou, China (201508020001). This work was supported in part by a grant from the BeiHao Stem Cell and Regenerative Medicine Translational Research Institute.

